# AMULETY: A Python package to embed adaptive immune receptor sequences

**DOI:** 10.1101/2025.03.21.644583

**Authors:** Meng Wang, Yuval Kluger, Steven H. Kleinstein, Gisela Gabernet

**Affiliations:** Program in Computational Biology and Biomedical Informatics, Yale University, New Haven, CT, USA; Department of Pathology, Yale School of Medicine, New Haven, CT, USA; Applied Mathematics Program, Yale University, New Haven, CT, USA; Department of Immunobiology, Yale School of Medicine, New Haven, CT, USA

**Keywords:** B cell receptor, embedding, machine learning, next generation sequencing, computational immunology

## Abstract

Large language models have been developed to capture relevant features of adaptive immune receptors, each with unique potential applications. However, the diversity in available models presents challenges in accessibility and usability for downstream applications. Here we present AMULETY (Adaptive imMUne receptor Language model Embedding Tool), a Python-based software package to generate language model embeddings for adaptive immune receptor sequences, enabling users to leverage the strengths of different models without the need for complex configuration. AMULETY offers functions for embedding adaptive immune receptor amino acid sequences using pre-trained protein or antibody language models for paired heavy and light chain or single chain sequences. We showcase the variability on the embedding space for several embeddings on a dataset of antibody binders to several SARS-CoV-2 epitopes and showed that different models may be effective at capturing different aspects of the distinctions between epitope groups. AMULETY is available under GPL v3 license from https://github.com/immcantation/amulety or via `pip` from the Python Package Index (PyPI) from https://pypi.org/project/amulety/.

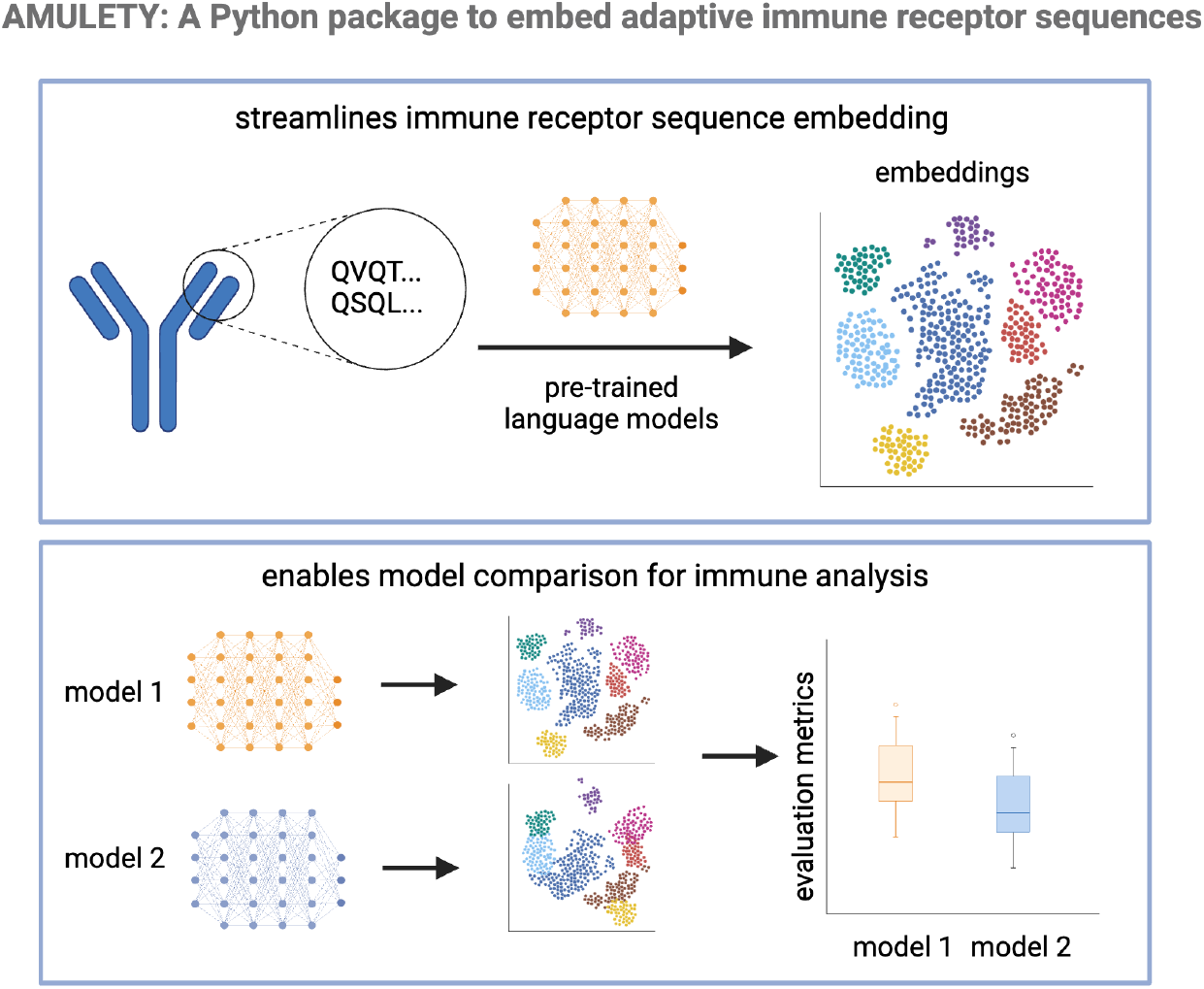

## 1. Introduction

The collection of B-cell receptors (BCRs), and T-cell receptors (TCRs) in an individual, also known as the adaptive immune receptor repertoire (AIRR), are key molecular components of the adaptive immune system that recognizes foreign and own antigens and initiate an immune response. They provide a record on current and past immune responses, and thus hold the potential to enable the prediction of the immune state of an individual, as well as inform on an individual’s response to challenges such as vaccines and immunotherapies.

Utilizing the AIRR sequences for these downstream applications often requires their translation into numerical representations that capture relevant properties of the sequences for specific prediction tasks, also termed embeddings. Several embedding methods have been described and shown to capture meaningful features of BCRs and TCRs relevant for various prediction applications, including some specifically trained on AIRR sequences such as immune2vec [1], AntiBERTy [2], AntiBERTa2 [3], BALM-paired [4]; as well as some trained on generic protein sequences, such as ESM2 [5].

Computing sequence representations with different embedding methods requires the installation of multiple library dependencies, as well as their specific configuration for usage on graphical processing units (GPU) infrastructure to speed up computation, which poses a technical challenge for inexperienced users. Here we present AMULETY, an open-source Python package to generate sequence embeddings for adaptive immune receptor sequences. The package facilitates the embedding computation of BCR and TCR sequences using different available methods with a simple command line interface, with support for GPU infrastructure. To ensure its interoperability with other AIRR data analysis tools, the AIRR rearrangement format is supported as an input. We also provide docker containers with the pre-installed tool dependencies for simplified installation and usage. AMULETY is available open-source on GitHub (https://github.com/immcantation/amulety) and freely distributed through the Python Package Index (PyPI) (https://pypi.org/project/amulety/) together with its usage documentation.

## 2. Implementation

### 2.1 Standardization of input sequences for embedding generation

AMULETY includes utility functions to standardize user-input sequences into amino acid sequences to generate pre-trained antibody language model embeddings. The input to this process is a data file in the Adaptive Immune Receptor Repertoire (AIRR) format [6], containing the nucleotide sequences and associated metadata. The function extracts the nucleotide sequences from the AIRR data file and formats them into a FASTA file, which serves as input to IgBLAST [7]. IgBlast is executed to perform sequence alignment to IMGT germline gene database and translation. The alignment and translations produced by IgBLAST include the full amino acid sequences and VDJ region translations.

The package supports embedding for both paired chain sequences as well as bulk sequencing data. For mixed datasets containing both single-cell and bulk entries, the tool filters and processes each type separately and combines the results together. Heavy-light chain pairing for cells with multiple light chains is achieved by identifying the most representative light chain sequences using consensus counts, with ties broken by selecting the first row encountered.

### 2.2 Pre-trained model embedding generation

To generate antibody sequence embeddings, multiple pre-trained language models were integrated into the package, each tailored for specific applications and architectural configurations (**Table 1**). Each embedding method runs on processed amino-acid sequences in AIRR-compliant format. For large sequence sets, the input sequences can be processed in mini-batches of customizable size, and GPU processing can be enabled for accelerated computation. Embedding results are provided as an aggregated output in formats such as .pt, .csv, or .tsv, depending on user requirements.

**Table 1.** Available antibody and protein language models for protein sequence embedding in the package.

| Model | Command | Embedding Dimension | Max Length | Notes |
| --- | --- | --- | --- | --- |
| AntiBERTy [2] | antiberty | 512 | 510 | Lightweight BERT model pre-trained on 588 million Observed Antibody Space (OAS) [8] heavy and light antibody sequences from multiple species. Suitable with low computational resources. |
| AntiBERTa2 [3] | antiberta2 | 1024 | 256 | RoFormer model pre-trained on 1.54 billion unpaired and 2.9 million paired human antibody sequences. Suitable for users needing a balance between computational cost and embedding dimension. |
| ESM2 (650M parameter) [5] | esm2 | 1280 | 512 | General protein language model pre-trained on 216 million UniRef50 [9] protein sequences. Ideal for complex tasks and requires heavy computational resources. |
| BALM-paired [4] | balm_paired | 1024 | 510 | RoBERTa-based large model pre-trained on 1.34 million concatenated heavy and light chain human antibody sequences. Specialized model for paired chain embeddings. |
| User-specified model | custommodel | Configurable | Configurable | Offers flexibility for users with pre-trained custom models. |

## 3. Example usage

We downloaded paired chain nucleotide sequences for antibodies binding to SARS-CoV-2 spike protein receptor binding domain including their epitope groups [10]. We standardized the nucleotide sequences into the protein sequences using the `amulety translate-igblast` command and generated the embeddings for individual cells in paired chain mode using all four models using `amulety <antiberty|antiberta2|balm_paired|esm2>`. The code for the two steps are shown below.

~~~
amulety translate-igblast <AIRR file> <output directory> <igblast data folder>
amulety <antiberty|antiberta2|esm2|balm_paired> --cell-id-col <cell_id_col>
--cache-dir <caching directory> --batch-size <mini batch size> <AIRR file>
<HL|H|L> <output embedding file>
~~~

We visualized the UMAP embeddings of the antibody language model embeddings and noticed differences in clustering of epitope groups between embedding models (**Figure 1A**). To formally quantify the differences of embedding distances in separating antibodies binding to different epitopes, we evaluated the embeddings using three metrics: Silhouette score, Davies-Bouldin index, and Calinski-Harabasz index. We found that the epitope-based clustering performance varied across models, with some embeddings exhibiting better separation of epitope groups than others (**Figure 1B**). The Silhouette scores measure how close antibodies are in the embedding space to others in the same epitope group, compared to antibodies in other epitope groups. Silhouette scores on the epitope groups are close to zero for all models, indicating weak epitope-based separation across the models. The Davies-Bouldin index focuses on the overall similarity between epitope groups and antiBERTa2 showed slightly better-defined antibody epitope clusters compared to other models. The Calinski-Harabasz index focuses on the ratio of between-cluster to within cluster dispersion and suggests that BALM-paired captured more distinct cluster structures. These results suggest that different antibody language models may be effective at capturing different aspects of the distinctions between epitope groups.

**Figure 1.**
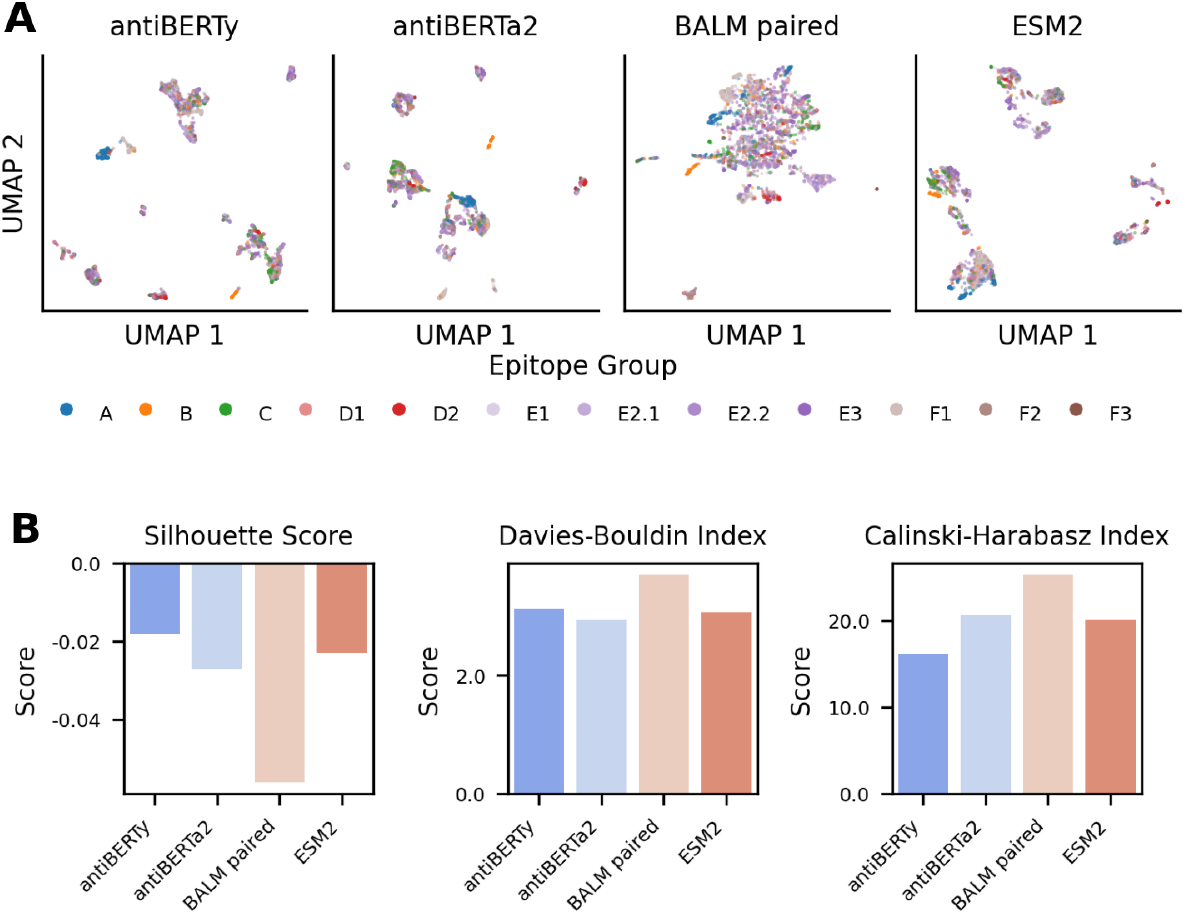
Comparison of Antibody Embedding Representations Using UMAP and Clustering Metrics. (A) UMAP projections of antibody embeddings generated by four different models: antiBERTy, antiBERTa2, BALM paired, and ESM2. Each column represents a UMAP projection for a specific embedding model, with points colored by their assigned epitope group. (B) Clustering evaluation metrics to assess the embedding models’ ability to separate antibodies binding to different epitopes, including Silhouette Score (higher is better), Davies-Bouldin Index (lower is better), and Calinski-Harabasz Index (higher is better).

## 4. Discussion

AMULETY is an open-source tool for generating language model embeddings of adaptive immune receptor sequences. By supporting multiple pre-trained models such as AntiBERTy, AntiBERTa2, BALM-paired, and ESM2, AMULETY allows users to leverage the unique strengths of each model for various tasks, depending on computational resources and the specific features of the immune repertoire they are studying.

A key extension for future developments is the inclusion of embedding methods specific for TCRs to allow for a more comprehensive analysis of adaptive immune repertoires. In addition, AMULETY is designed to be modular and integrate seamlessly with other AIRR processing tools that support the AIRR format, including the Immcantation framework, which supports other AIRR data analysis tools. Its modularity ensures that the tool can be expanded to accommodate future developments in both immune repertoire analysis and machine learning models.

In conclusion, AMULETY provides a user-friendly command line interface to access pre-trained antibody language models for sequence embedding and reduces the technical barriers often associated with using advanced machine learning models to extract meaningful insights from adaptive immune receptor sequences. This ensures that researchers can focus on the biological interpretation of their data rather than the complexities of model configuration. The ability to use the tool with both CPU and GPU support allows for flexibility in resource usage, making it accessible to a wider range of users, from those with limited computational resources to those with access to high-performance computing environments.

## CRediT authorship contribution statement

**MW**: Methodology, Software, Formal analysis, Validation, Data curation, Writing - original draft. **YK**: Conceptualization, Funding acquisition, Supervision, Project administration, Writing - review & editing. **SHK**: Conceptualization, Funding acquisition, Supervision, Project administration, Writing - review & editing. **GG**: Software, Formal analysis, Funding acquisition, Supervision, Validation, Writing - original draft.

## Acknowledgements

The authors thank Edel Aron for helpful feedback on the package during its testing.

## Supplementary material

Supplementary material associated with this article can be found at the online version.

